# Tissue-distinct Features of Follicular Cytotoxic CD8^+^ T Cells in *Trypanosoma cruzi* infection

**DOI:** 10.64898/2026.02.25.707971

**Authors:** Yamila Gazzoni, Laura Almada, Julio C. Gareca, Cinthia C. Stempin, Carolina L. Montes, Eva V. Acosta-Rodríguez, Adriana Gruppi

**Affiliations:** Departamento de Bioquímica Clínica, Facultad de Ciencias Químicas (FCQ), Universidad Nacional de Córdoba (UNC), Córdoba, Argentina; Centro de Investigaciones en Bioquímica Clínica e Inmunología (CIBICI - CONICET), Córdoba, Argentina

**Keywords:** Follicular cytotoxic CD8^+^T cells - CD8^+^ T cell - *Trypanosoma cruzi* - Chagas disease

## Abstract

T follicular cytotoxic (Tfc) cells are a specialized subset of CD8⁺ T lymphocytes that combine helper and cytotoxic functions. While their phenotypic and functional properties have been described in several infectious and inflammatory settings, little is known whether Tfc cells exhibit distinct characteristics across secondary lymphoid organs. Using acute *Trypanosoma cruzi* infection as a model of systemic immune activation, we performed a comparative phenotypic, transcriptomic, metabolic, and functional characterization of Tfc cells arising in the spleen and inguinal lymph nodes. Tfc cells emerged transiently in both organs with similar kinetics and antigen specificity yet segregated primarily according to tissue of analysis at the transcriptional level. Splenic Tfc cells exhibited enhanced glycolytic and mTORC1-associated signatures, increased mitochondrial mass, and stronger cytotoxic and B cell helper functions, promoting immunoglobulin production and plasmablast death. In contrast, Tfc cells from lymph node preferentially displayed memory-associated features, including higher CD127 and CD122 expression, increased IL-2 production, and preferential persistence during chronic infection. Together, these findings demonstrate that Tfc cells are not a homogeneous population but can adopt organ-associated transcriptional, metabolic, and functional states during *T. cruzi* infection.

## Introduction

T follicular cytotoxic (Tfc) cells constitute a specialized subset of CD8⁺ T lymphocytes that integrate hallmark features of CD4⁺ T follicular helper (Tfh) cells with the canonical cytotoxic program of CD8⁺ T cells (1,2). Similar to Tfh cells, Tfc cells express CXCR5, PD-1, ICOS, and the transcription factor Bcl6, which supports their localization within B-cell follicles or adjacent perifollicular niches and enables functional interactions with B cells (3,4). In contrast to classical Tfh cells, Tfc cells retain a potent cytotoxic profile, characterized by Granzyme B and Perforin expression and gene signatures associated with CD8⁺ T cell–mediated killing (4).

We previously identified a Tfc population that emerges in the spleen of *T. cruzi*-infected mice during the acute phase of infection and is no longer detectable thereafter (5). In this setting, Tfc cells peak in parallel with plasmablast expansion, and arise prior to germinal center formation. Splenic Tfc cells display elevated expression of molecules associated with B cell help, inflammatory chemokine receptors, and transcription factors linked to effector activity. *In vitro* analyses showed that Tfc cells modulate the humoral response in *T. cruzi*–infected mice, integrating effector, helper, and regulatory programs (5).

Secondary lymphoid organs (SLO) are anatomically and functionally specialized sites where adaptive immune responses are initiated and organized (6,7). In addition to immune cells, SLO contain structural stromal networks and vascular compartments that regulate antigen delivery, cellular positioning, and lymphocyte trafficking (8,9). Although spleen and lymph nodes (LN) share core organizational principles, they differ in antigen access and activation dynamics. LN drain peripheral tissues and are primarily exposed to lymph-borne antigens, whereas the spleen filters blood and is specialized for responses to circulating pathogens (10). During an infection, these organs undergo dynamic changes in cellular composition and immune activation states, driven by inflammatory cues and pathogen dissemination (11). In *T. cruzi* infection, both spleen and LN exhibit marked alterations in immune cell distribution and differentiation programs during the acute phase (12–16).

Whether Tfc cells display comparable phenotypic and transcriptional programs across distinct SLO during acute infection remains unknown. Defining potential organ-associated differences is important to understand how immune responses are spatially organized and diversified within the same host. Here, we performed an in-depth phenotypic and transcriptomic characterization of Tfc cells emerging during the acute phase of *T. cruzi* infection in the spleen and inguinal LN, to determine whether tissue environments are associated with specific characteristics on this specialized CD8^+^ T cell subset.

## Materials and Methods

### Mice and infection

Female wild-type C57BL/6 mice (8–12 weeks old) were housed under controlled temperature and humidity with a 12-hour light/dark cycle. The colony, originally obtained from the Facultad de Veterinaria, Universidad Nacional de La Plata, has been maintained at the CIBICI-CONICET, FCQ-UNC Animal Facility for over seven years.

To induce *T. cruzi* infection, mice were injected intraperitoneally with 5×10³ trypomastigotes of the Tulahuén strain of *T. cruzi* in 0.2 mL PBS. Age-matched controls received PBS alone.

### Cell suspensions

For this study, spleen and inguinal LN were selected because they undergo marked hypertrophy during the acute phase of *T. cruzi* infection (13), instead of mesenteric LN which suffer marked atrophy (16). Tissues were harvested, mechanically dissociated, erythrocytes were lysed with Tris–ammonium chloride buffer, and viable cells were counted by trypan blue exclusion using a hemocytometer.

### Antibodies and flow cytometry

Surface and intracellular staining: cells were stained with fluorochrome-conjugated Abs and a fixable viability dye for 25 min at 4°C. *T. cruzi-*specific CD8^+^T cells were detected using H-2K(b)/TSKB20 tetramers. CXCR5 or GLUT-1 were revealed with biotinylated Abs followed by streptavidin. Intracellular transcription factors and cytokines were stained after fixation/permeabilization using Foxp3 or Cytofix/Cytoperm buffers.

Metabolic and signaling staining: Mitochondrial mass, membrane potential, and mROS were assessed with MitoTracker Green/Orange or MitoStatus Red and MitoSOX, respectively. For phosphorylated protein, cells were stained for surface markers and subsequently fixed and permeabilized according to the target phosphoprotein. Detection of phospho-mTOR (Ser2448) and phospho-AKT (Ser473) was performed using the Foxp3 Staining Buffer Set, whereas phospho-p70 S6K (Thr389) was analyzed following fixation with 4% paraformaldehyde and permeabilization with BD Phosflow™ Perm Buffer. Phospho-p70-S6K was detected using an AlexaFluor594–conjugated secondary Ab.

Data were collected on LSRFortessa X-20 flow cytometer (BD Biosciences). Abs and reagents are listed in Table S1.

### Cell sorting

Spleen and inguinal LN cell suspensions were stained with fluorochrome-conjugated Abs and a fixable viability dye and sorted on a FACSAria II (BD). Live lymphocytes were gated and Tfc (CXCR5^+^PD-1^+^CD8^+^CD4^-^B220^-^), non-Tfc (CXCR5^-^PD-1^-^CD8^+^CD4^-^B220^-^), naïve B cells (B220^+^IgD^+^), and plasmablasts (B220^int^CD138^+^IgD^-^) were purified as described (5).

### RNA sequencing and analysis

Tfc and non-Tfc cells were sorted from spleen and inguinal LN of *T. cruzi*–infected mice at 18 dpi. RNA isolation, library preparation, and sequencing were performed as previously described (5). In brief, reads were quality filtered, aligned to the *Mus musculus* genome (GRCm38), and differential expression was analyzed using DESeq2 with Wald statistics. Genes with adjusted *p*<0.05 and log_2_fold change>1 were considered differentially expressed.

### *In vitro* co-cultures

Co-cultures were performed as described previously (5). In brief, sorted naïve B cells or plasmablasts were co-cultured with Tfc cells in anti-CD3/anti-CD28–coated plates for 20 h. Supernatants were collected for immunoglobulin analysis. Plasmablast survival was evaluated using a caspase-3/7 assay; live cells were defined as Caspase-3^-^SYTOX^-^CD8^-^B220^int^. Killing was calculated as: % killing = 1−(%live PB+Tfc / %live PBalone)×100.

### *In vitro* functional stimulation assay

Spleen or inguinal LN cell suspensions were stimulated with the TSKB20 peptide (5 μg/mL, ANYKFTLV), or with PMA and ionomycin (1 μg/mL), in the presence of Brefeldin A and Monensin (5). Anti-CD107a Ab was included throughout the incubation period. Following culture, cells were processed for flow cytometry analysis. Data acquisition was performed using a Cytek Northern Light flow cytometer (Gematec).

For cytokine analysis, sorted Tfc cells from both SLO were stimulated for 4 h with anti-CD3 (0.5 μg/mL) plus anti-CD28 (0.2 μg/mL) coated 96-well U-bottom plates.

### Immunoglobulin and cytokine quantification

Immunoglobulin isotype and cytokine concentrations were quantified in co-culture supernatants by LEGENDplex (5). Data were acquired on a FACSCanto II flow cytometer and analyzed using LEGENDplex Data Analysis Software. For statistical analyses, ND values were imputed using the lower limit of detection of the assay.

### Glucose uptake assay

Glycolytic activity was assessed using a 2-NBDG uptake assay (17). Splenic and inguinal LN cell suspensions obtained at 18 dpi from infected mice were incubated with 2-NBDG, washed, and subsequently stained for surface markers. Mean fluorescence intensity (MFI) of 2-NBDG uptake was then quantified by flow cytometry.

### Statistical analysis and data visualization

Statistical analyses were performed using GraphPad Prism (v8.0.1). Data normality was assessed using the Shapiro–Wilk test. Normally distributed data were analyzed using Student’s *t*-test or one-way ANOVA with Bonferroni’s post-hoc test, whereas non-normally distributed data were analyzed using the Mann–Whitney U or Kruskal–Wallis test, as appropriate. *P* ≤ 0.05 was considered statistically significant. Outliers were identified using the ROUT method. Data are presented as mean ± standard deviation (SD), and sample size is indicated in figure legends. Flow cytometry data were analyzed using FlowJo (vX.0.7 and 10.8.1) with gating defined using negative and FMO controls.

Transcriptomic visualization and pathway enrichment analyses were performed using the SRplot (18) and iDEP online platform (19). Gene Set Enrichment Analyses (GSEA) was conducted using the GSEA software and gene sets from the Molecular Signatures Database (MSigDB) (20).

### AI language model assistance

ChatGPT (OpenAI) assisted with language editing for clarity and grammar; all text was critically reviewed and revised by the authors.

## Results

### Splenic and LN-Tfc cells exhibit similar kinetic but different transcriptional profiles

To evaluate Tfc cell characteristics in different SLO, we first analyzed the kinetics of these cells in the spleen (S-Tfc) and inguinal LN (LN-Tfc) from *T. cruzi*-infected mice. Tfc cells were identified by flow cytometry based on CXCR5 and PD-1 expression on CD4^-^B220^-^CD8^+^ cells (Figure 1A). Consistent with our previous observations (5), Tfc cells were absent in non-infected mice but emerged transiently during the acute phase of *T. cruzi* infection, peaking at 18 dpi with comparable kinetics in spleen and LN (Figure 1A).

**Figure 1.**
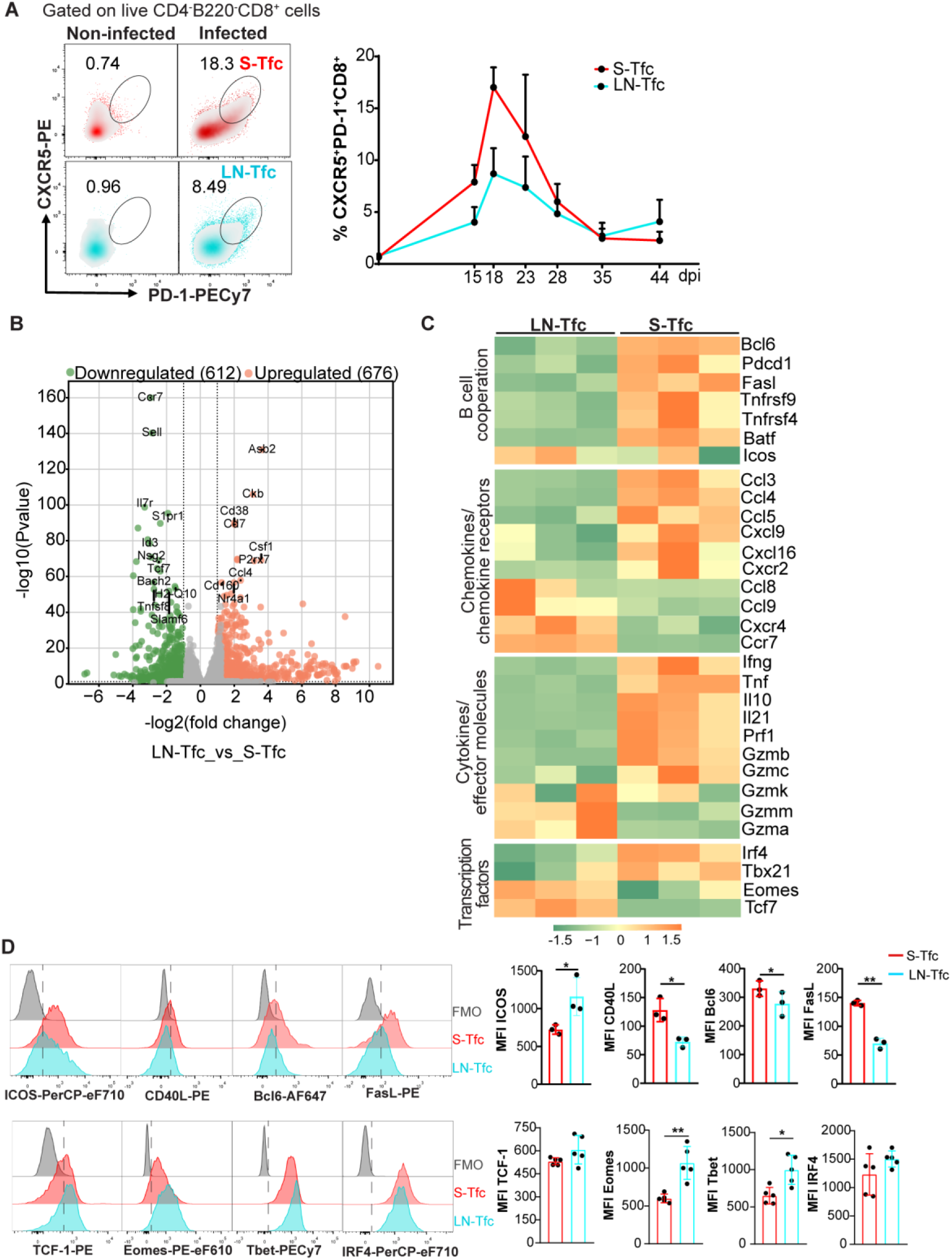
Dynamic, transcriptional and phenotypic features of Tfc cells from SLO during acute phase of *T. cruzi* infection. C57BL/6 mice were intraperitoneally injected with PBS (non-infected) or with 5000 trypomastigotes of *T. cruzi* Tulahuén strain (infected). (A) Tfc cells from spleen (S-Tfc, red) and inguinal lymph node (LN-Tfc, cyan) were analyzed by flow cytometry at different days post-infection (dpi). Representative density plots and kinetics showing the frequency of Tfc cells (CXCR5^+^PD-1^+^CD8^+^) gated on live CD4^-^B220^-^CD8^+^ T lymphocytes in non-infected and at 18 dpi infected mice. (B) S-Tfc and LN-Tfc cells were isolated by cell sorting from *T. cruzi*-infected mice at 18 dpi and subjected to bulk RNA sequencing. Volcano plot displaying differentially expressed genes between S-Tfc and LN-Tfc cells. Significantly downregulated (green dots) and upregulated (orange dots) genes were defined by log2fold change>1 and adjusted p-value < 0.05. Selected genes are labeled. (C) Heatmap showing the relative expression of selected genes in S-Tfc and LN-Tfc cells. The color scale represents the Z-score of gene expression, with green indicating lower expression and orange higher expression. (D) Representative histograms and statistical analysis of mean fluorescence intensity (MFI) of molecules associated with B cell help (ICOS, CD40L and FasL) and Tfh program (Bcl6), and transcription factors (TCF-1, Eomes, Tbet and IRF4) in Tfc cells from both SLO. (A, D) Data are presented as mean ± SD; N = 3-5 mice and are representative from 3 independent experiments. Statistical significance was determined by a paired *t*-test (*p < 0.05, **p < 0.01). (B, C) Each biological replicate (N = 3) consisted of pooled cells from three mice.

To examine the transcriptional profiles of Tfc cells across SLO, we performed comparative RNA-seq analysis of S-Tfc and LN-Tfc cells isolated by cell sorting from 18 dpi–infected mice. To determine whether potential transcriptional differences reflected Tfc cell–intrinsic features or broader tissue-associated effects on CD8^+^ T cells, non-follicular CD8^+^ T cells (non-Tfc, CXCR5^-^PD-1^-^CD8^+^) from each organ were also included in the Principal Component Analysis. Both Tfc and non-Tfc subsets segregated primarily according to tissue origin along PC1, which accounted for 57% of the total variance (Figure S1), indicating a dominant effect of tissue environment on CD8^+^ T cell transcriptional programs.

Focusing specifically on the Tfc cell population, we observed differential gene expression between S-Tfc and LN-Tfc cells. A total of 1,288 transcripts were differentially expressed between the two Tfc populations, with 676 genes upregulated and 612 genes downregulated in S-Tfc cells relative to LN-Tfc cells (Figure 1B). Visualization of selected gene sets highlighted these differences, showing differential expression patterns between S-Tfc and LN-Tfc cells (Figure 1C). S-Tfc cells were enriched in transcripts associated with B cell cooperation and interaction, including the transcription factors *Bcl6* and *Batf*, as well as effector and costimulatory molecules such as *Pdcd1*, *Tnfrsf9*, *Tnfrsf4*, and *Fasl*. Moreover, S-Tfc cells showed increased expression of transcripts encoding inflammatory chemokines such as *Ccl3*, *Ccl4*, *Ccl5*, *Cxcl9*, *Cxcl16*, and the chemokine receptor *Cxcr2*, together with elevated expression of transcripts encoding effector and immunomodulatory molecules, including *Ifng*, *Tnf*, *Il10, Il21, Prf1, Gzmb*, and *Gzmc*, while exhibiting reduced expression of transcripts associated with lymphoid homing receptors, namely *Ccr7* and *Cxcr4*, compared with LN-Tfc cells. S-Tfc cells were further enriched in transcripts encoding transcription factors linked to CD8^+^ T cell activation and effector differentiation, such as *Irf4* and *Tbx21* (21). In contrast, LN-Tfc cells preferentially expressed transcripts associated with memory-related programs, notably *Tcf7* and *Eomes* (22), and displayed higher expression of *Gzma* and *Gzmm*.

To validate the RNA-seq data, we assessed selected proteins by flow cytometry. S-Tfc cells displayed higher expression of the B cell help–associated molecules CD40L and FasL, together with increased levels of the transcription factor Bcl6, whereas LN-Tfc cells exhibited increased expression of ICOS, as determined by mean fluorescence intensity (MFI). In addition, LN-Tfc cells showed higher protein levels of the transcription factors Eomes and T-bet (Figure 1D), despite the enrichment of *Tbx21* transcripts in S-Tfc cells.

[ufig1]

### S-Tfc and LN-Tfc cells differ in metabolic and mitochondrial features

Given the marked transcriptional divergence between S-Tfc and LN-Tfc cells, and the established role of metabolic programs in CD8^+^ T cell biology (23), we next examined whether the transcriptional differences observed between S-Tfc and LN-Tfc cells were accompanied by distinct metabolic features. Gene set enrichment analysis (GSEA) of RNA-seq data revealed a significant enrichment of *Glycolysis* and *mTORC1 signaling* gene sets in S-Tfc cells compared with LN-Tfc cells (Figure 2A).

**Figure 2.**
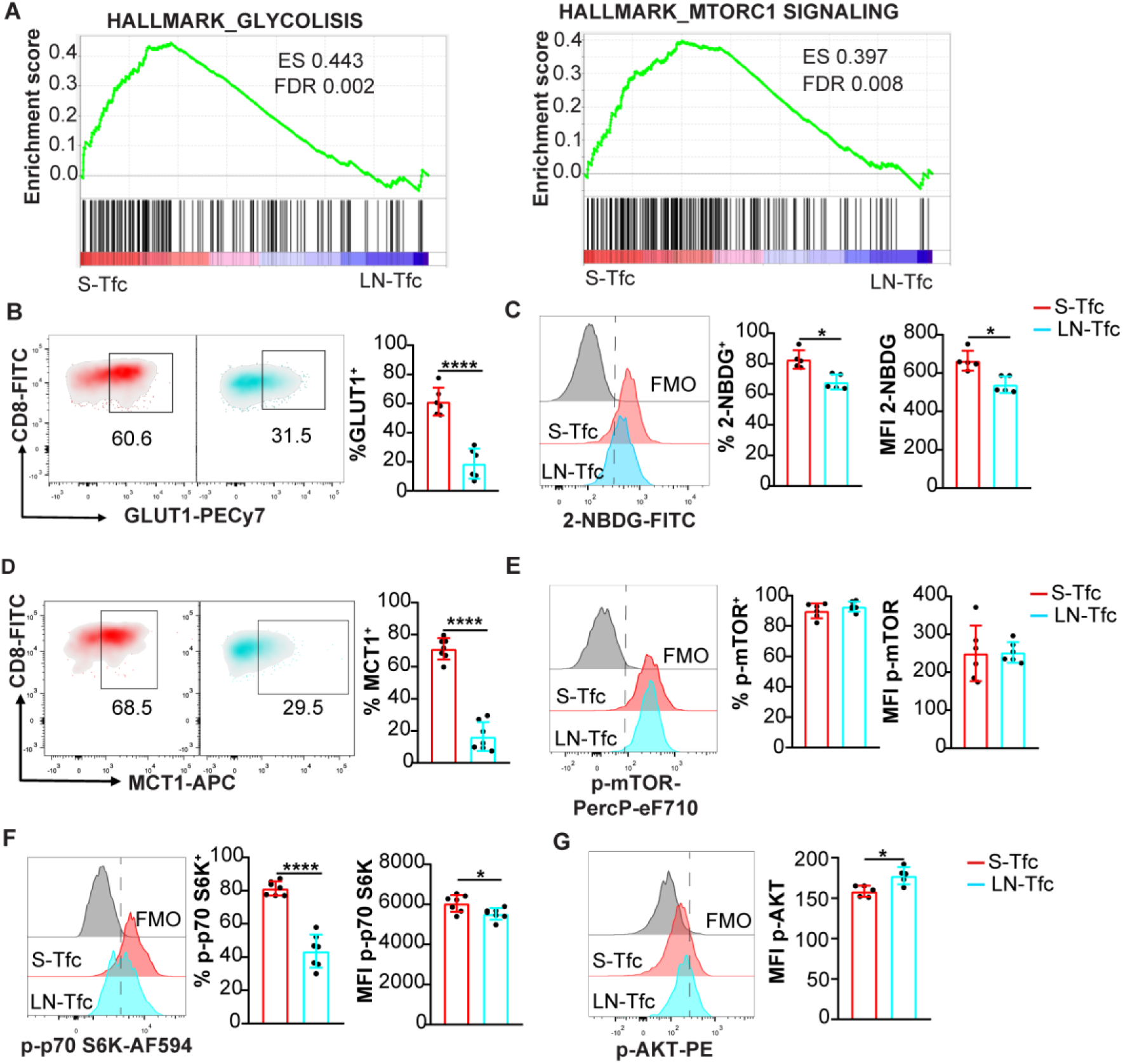
Metabolic profiling of Tfc cells in SLO. (A) Gene Set Enrichment Analysis (GSEA) of bulk RNA-seq data from sorted S-Tfc and inguinal LN-Tfc cells obtained at 18 dpi, highlighting enrichment of *glycolysis* and *mTORC1 signaling* pathways using the MSigDB Hallmark (MH) mouse collection. ES, enrichment score; FDR, false discovery rate. Genes enriched in S-Tfc cells are located toward the left of the ranked list, whereas genes preferentially expressed in LN-Tfc cells are located toward the right. (B–G) Spleen and inguinal LN cells were obtained from *T. cruzi*–infected mice at 18 dpi and analyzed ex vivo by flow cytometry. (B, D) Representative density plots and quantification of the frequency of (B) GLUT1^+^ and (D) MCT1^+^ cells among gated S-Tfc and LN-Tfc populations. (C, E, F) Representative histograms and quantification of both the frequency of positive cells and MFI for (C) 2-NBDG uptake, (E) phosphorylated mTOR (p-mTOR), and (F) phosphorylated p70 S6 kinase (p-p70 S6K) in S-Tfc and LN-Tfc cells. (G) Representative histograms and quantification of phosphorylated AKT (p-AKT) MFI in S-Tfc and LN-Tfc cells. Data are presented as mean ± SD; *N* = 5–7 mice. (B–G) Data are representative of three independent experiments. Statistical significance was determined using an unpaired two-tailed *t*- test (*p*\*<0.05, *p*\*\*\*<0.001, *p*\*\*\*\*<0.0001).

To validate these signatures at the protein level, we assessed metabolic parameters by flow cytometry. At 18 dpi, S-Tfc cells exhibited a higher frequency of GLUT1^+^ cells (Figure 2B) and enhanced glucose uptake, reflected by both a higher frequency of 2-NBDG^+^ cells and an increased 2-NBDG MFI, compared with LN-Tfc cells (Figure 2C). Consistent with enhanced glucose utilization, S-Tfc cells displayed a higher frequency of cells expressing MCT1, a transporter involved in the transmembrane transport of lactate and pyruvate (24) (Figure 2D). No significant differences were observed in either the frequency or the MFI of cells expressing phosphorylated-mTOR (p-mTOR) (Figure 2E). However, phosphorylated-p70 S6 kinase (p-p70 S6K), a downstream readout of mTORC1 activity (25), was significantly increased in S-Tfc cells reflected by both a higher frequency of p-p70 S6K^+^ cells and an increased p-p70 S6K MFI (Figure 2F). In contrast, LN-Tfc cells exhibited higher levels of phosphorylated-AKT (p-AKT, Figure 2G), consistent with mTORC2-associated signaling (25).

In addition to glycolytic activity, mitochondria play a key role in CD8^+^ T cell activation and function (26). We therefore next assessed mitochondrial parameters in S-Tfc and LN-Tfc cells. S-Tfc cells displayed a higher frequency of cells double-positive for MitoTracker Green (MTGreen) and MitoTracker Orange (MTOrange), indicating a greater proportion of mitochondria with preserved membrane potential (Figure 3A). Although no differences were observed in MTOrange MFI, S-Tfc cells exhibited higher levels of MTGreen MFI (Figure 3B), suggestive of higher mitochondrial mass. Finally, MitoSOX staining demonstrated a significantly higher frequency of mROS⁺ cells in S-Tfc cells compared with LN-Tfc cells (Figure 3C), consistent with the role of mitochondrial mROS as signaling mediators in T cell activation and differentiation (27).

**Figure 3.**
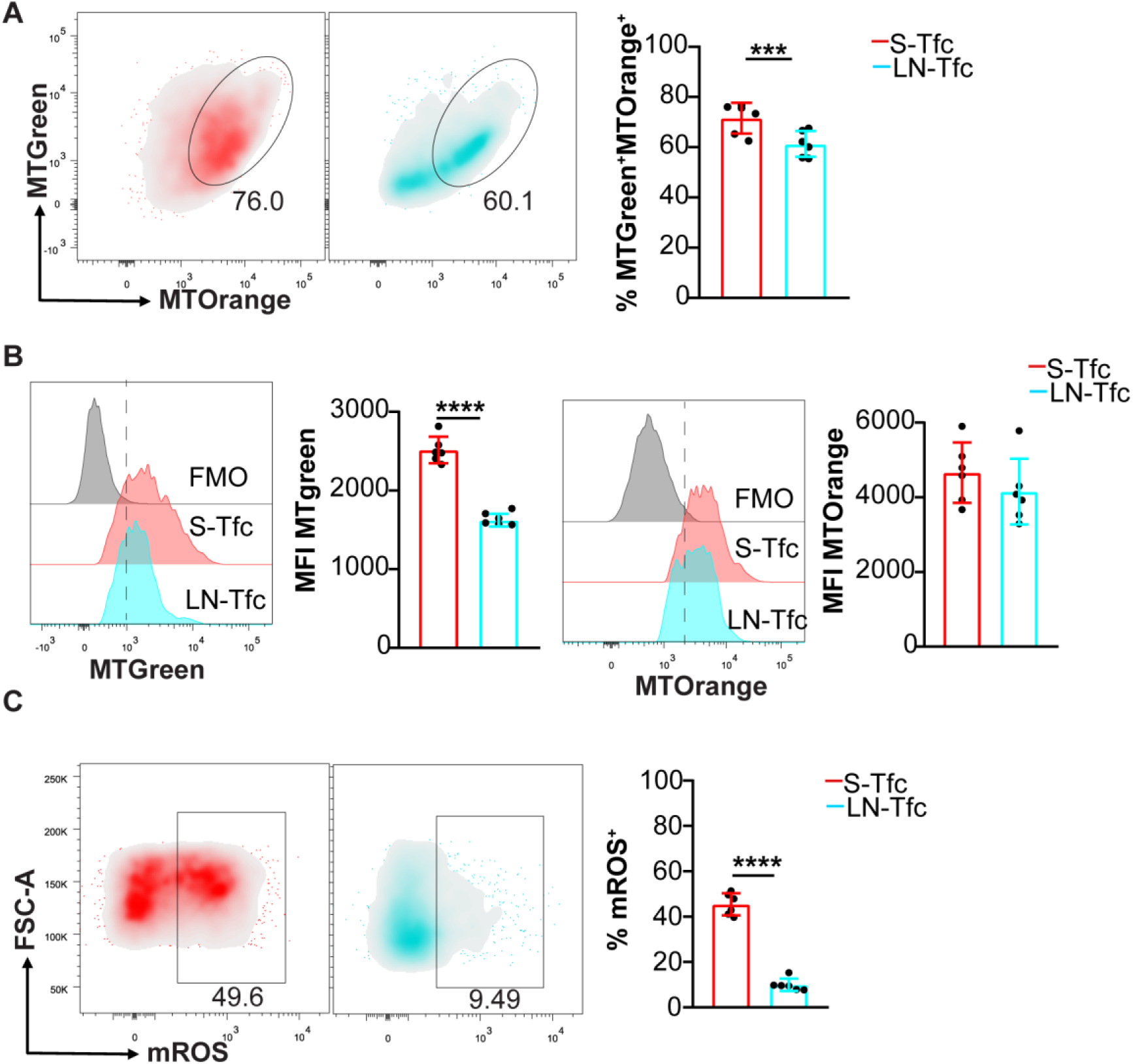
Mitochondrial features analysis on S-Tfc and LN-Tfc cells. Cells from spleen and inguinal LN were obtained at 18 dpi from *T. cruzi* infected mice and the expression of mitochondria-associated markers was evaluated by flow cytometry. (A) Representative density plots and statistical analysis of frequency of MitoTracker Green^+^MitoTracker Orange^+^ (MTGreen^+^MTorange^+^) cells among gated Tfc cells. (B) Representative histograms and statistical analysis of MFI of MTGreen and MTorange on MTGreen^+^MTorange^+^ Tfc cells. (C) Representative density plots and statistical analysis of frequency of mROS^+^ gated on Tfc cells. (A-C) Data are presented as mean ± SD; *N* = 6–7 mice and are representative of two (C) and three (A-B) independent experiments. Statistical significance was determined using an unpaired two-tailed *t*- test (*p*\*\*\*<0.001, *p*\*\*\*\*<0.0001).

### Splenic and LN-derived Tfc cells display distinct differentiation profiles and persistence patterns

Given the distinct metabolic profiles identified in S-Tfc and LN-Tfc cells, we next examined whether these subsets also differed in their differentiation states. We analyzed the distribution of effector and memory CD8^+^ T cell subsets within the Tfc compartment in spleen and inguinal LN at 18 dpi. Flow cytometric analysis revealed that the majority of S-Tfc cells exhibited an effector/effector memory phenotype (Eff, Em, CD44^+^CD62L^-^), whereas LN-Tfc cells were composed of comparable proportions of Eff/Em and central memory cells (CM, CD44^+^CD62L^+^) (Figure 4A). Further analysis based on KLRG1 and CD127 expression showed a higher frequency of short-lived effector cells (SLEC, KLRG1^+^CD127^-^) and early effector cells (EEC, KLRG1^-^CD127^-^) within the S-Tfc compartment compared with LN-Tfc cells. In contrast, LN-Tfc cells displayed a higher proportion of memory precursor effector cells (MPEC, CD127^+^KLRG1^-^) than S-Tfc cells (Figure 4B). Consistent with the increased representation of memory-associated subsets within LN-Tfc cells, these cells also exhibited higher expression of CD122 (28) compared with S-Tfc cells (Figure 4C).

**Figure 4.**
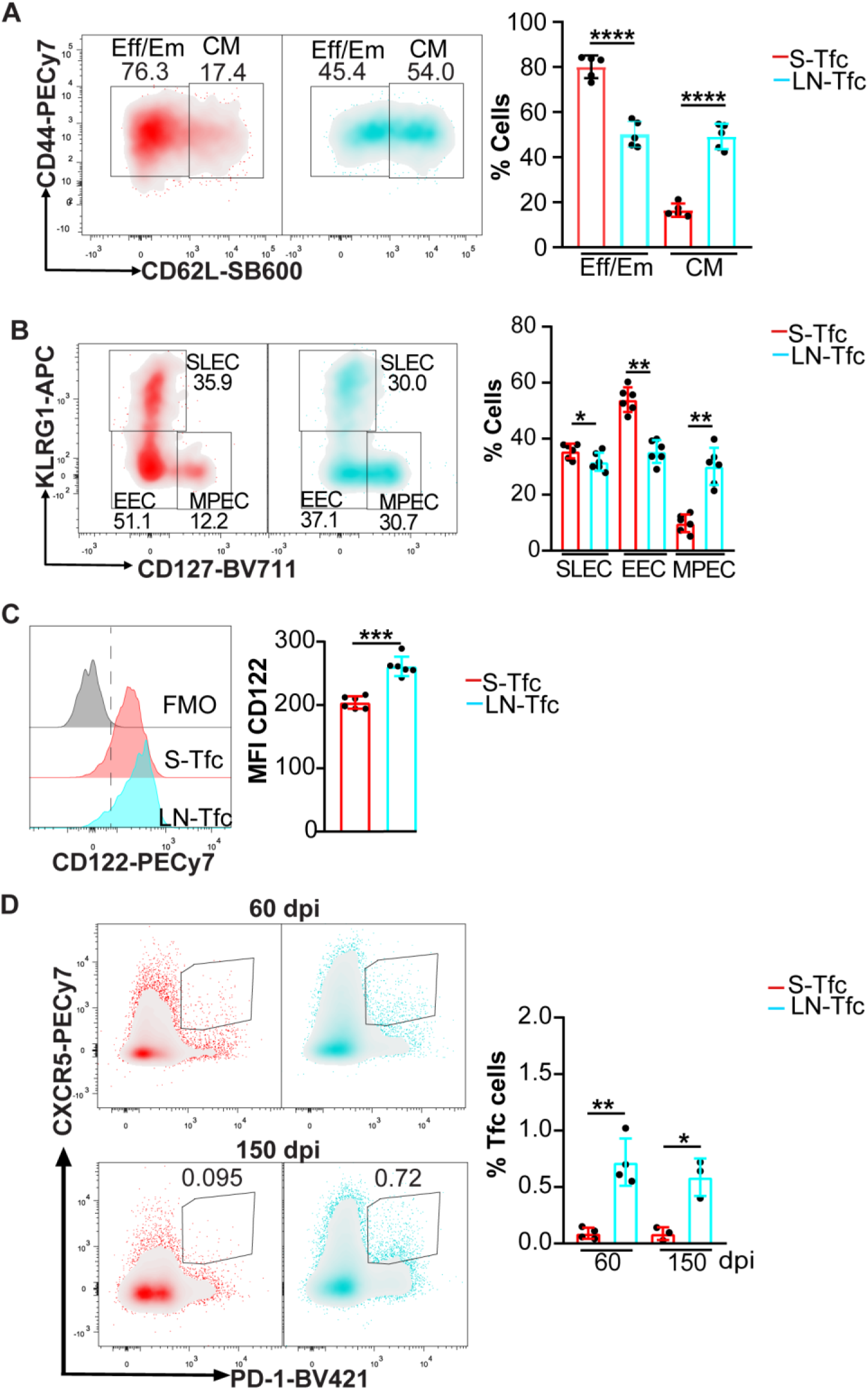
Differentiation state of splenic and LN-derived Tfc cell populations. Spleen and inguinal LN cells were obtained from *T. cruzi*–infected mice at 18 dpi (A–C) or at 60 and 150 dpi (D) and analyzed by flow cytometry. (A-B) Representative density plots and statistical analysis of the frequency of (A) effector/effector memory (Eff/Em, CD44^+^CD62L^-^) and central memory (CM, CD44^+^CD62L^+^) cells; and (B) short-lived effector cells (SLEC, KLRG1^+^CD127^-^), early effector cells (EEC, KLRG1^-^CD127^-^) and memory precursors effector cells (MPEC, KLRG1^-^CD127^+^) among gated S-Tfc and LN-Tfc cells at 18 dpi. (C) Representative histograms and statistical analysis of CD122 MFI on Tfc cells from both SLO at 18 dpi. (D) Representative density plots and statistical analysis of the frequency of Tfc (CXCR5^+^PD-1^+^CD8^+^) cells gated on CD8^+^CD4^-^B220^-^ cells from spleen and inguinal LN at 60 and 150 dpi. Data are presented as mean ± SD; *N* = 3-6 mice. (A–C) Data are representative of three independent experiments, and (D) data are representative of two independent experiments. Statistical significance was determined using an unpaired two-tailed *t*-test (*p < 0.05, **p < 0.01, ***p < 0.001).

Given the enrichment of memory precursor populations within LN-Tfc cells at 18 dpi, we next assessed whether Tfc cells were maintained at later time points (60 and 150 dpi) in a tissue-dependent manner. At both time points, flow cytometric analysis revealed that Tfc (CXCR5^+^PD-1^+^CD8^+^CD4^-^B220^-^) cells were almost absent from the spleen, whereas a detectable proportion of Tfc cells persisted in the inguinal LN (Figure 4D), indicating differential persistence of Tfc cells across SLO.

### Splenic and LN–Tfc cells exhibited distinct effector molecule and cytokine production profiles

Having identified distinct transcriptional, metabolic, and differentiation profiles between S-Tfc and LN-Tfc cells, we next examined their parasite antigen specificity, effector molecule expression, and cytokine production. We first quantified the frequency of *T. cruzi*–specific Tfc cells using the immunodominant CD8^+^ T cell *T. cruzi* epitope TSKB20 (29). At 18 dpi, no differences were detected in the frequency of TSKB20-specific Tfc cells between SLO (Figure 5A).

**Figure 5.**
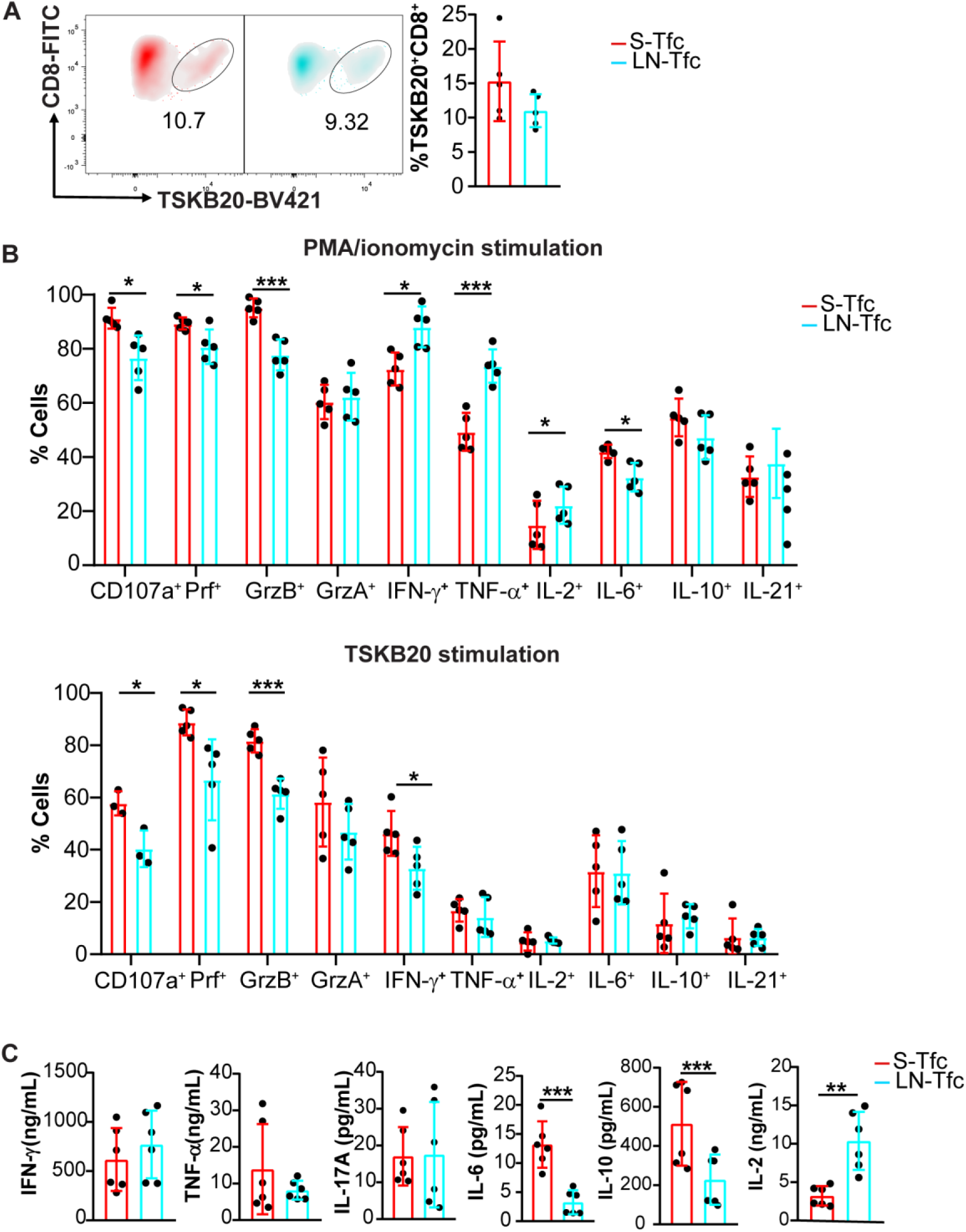
Cytokine production profiles and antigen specificity of Tfc from spleen and LN. (A, B) Spleens and inguinal LN were collected from *T. cruzi* infected mice at 18 dpi and cells were analyzed by flow cytometry. (A) Representative density plots and statistical analysis of the frequency of TSKB20^+^CD8^+^ cells among gated S-Tfc and LN-Tfc cells. (B) Statistical analysis of the frequency of CD107a^+^ Tfc cells and effector molecules-producing cells (Perforin^+^, Granzyme B^+^, Granzyme A^+^, IFN-γ^+^, TNF-α^+^, IL-2^+^, IL-6^+^, IL-10^+^, and IL-21^+^) following 4 h stimulation with either PMA/ionomycin (upper panel) or the TSKB20 peptide (lower panel) among gated S-Tfc and LN-Tfc cells. (C) Statistical analysis of concentration of IFN-γ, TNF-α, IL-17A, IL-6, IL-10, and IL-2, in culture supernatants from sorted S-Tfc and LN-Tfc cells stimulated with anti-CD3 plus anti-CD28 for 4 h. (A, B) Data are representative from two independent experiments (N= 5 mice). (C) Data are pooled from two independent experiments (N = 6-7 mice total). Data are presented as mean ± SD. Statistical significance was determined using an unpaired two-tailed *t*-test (*p*\*<0.05, *p*\*\*< 0.01, *p*\*\*\*<0.001). Abbreviations: Prf, perforin; GrzB, granzyme B; GrzA, granzyme A.

Cells isolated from the spleen and inguinal LN at 18 dpi were stimulated *in vitro* with either PMA/ionomycin or TSKB20, followed by flow cytometric analysis of cytotoxic mediators and cytokine-producing cells. Under both stimulation conditions, S-Tfc cells exhibited significantly higher frequencies of CD107a^+^, perforin^+^ (Prf^+^), and granzyme B^+^ (GrzB^+^) cells compared with LN-Tfc cells, suggesting an enhanced cytotoxic effector potential (Figure 5B). Following PMA/ionomycin stimulation, S-Tfc cells displayed increased frequencies of IL-6–producing cells, whereas LN-Tfc cells showed higher frequencies of IFN-γ^+^, TNF-α^+^, and IL-2^+^ cells. In contrast, after TSKB20 stimulation, S-Tfc cells showed a higher frequency of IFN-γ–producing cells, while no differences were observed in the frequency of TNF-α⁺, IL-2⁺, IL-6⁺, IL-10⁺, or IL-21⁺ cells between S-Tfc and LN-Tfc populations. No differences were detected in Granzyme A (GrzA) expression under either stimulation condition.

To further assess cytokine secretion, purified Tfc cells from both SLO were stimulated with anti-CD3 and anti-CD28 for 4 h, and cytokine concentrations in culture supernatants were quantified (Figure 5C). Under these conditions, S-Tfc and LN-Tfc cells secreted comparable levels of IFN-γ, TNF-α, and IL-17A. However, S-Tfc cells produced significantly higher amounts of IL-10 and IL-6, whereas LN-Tfc cells secreted higher levels of IL-2.

### Tfc cells from distinct SLO differentially modulate B cell responses

In light of our previous findings that S-Tfc cells promote Ig production and induce plasmablast death (5), and the functional differences identified here, we next examined whether S-Tfc and LN-Tfc cells differed in their capacity to modulate B cell responses. In line with this, GSEA on RNA-seq data uncovered a significant enrichment in S-Tfc cells of gene sets associated with *T cell–mediated cytotoxicity* and *humoral immune response mediated by circulating immunoglobulin* (Figure 6A).

**Figure 6.**
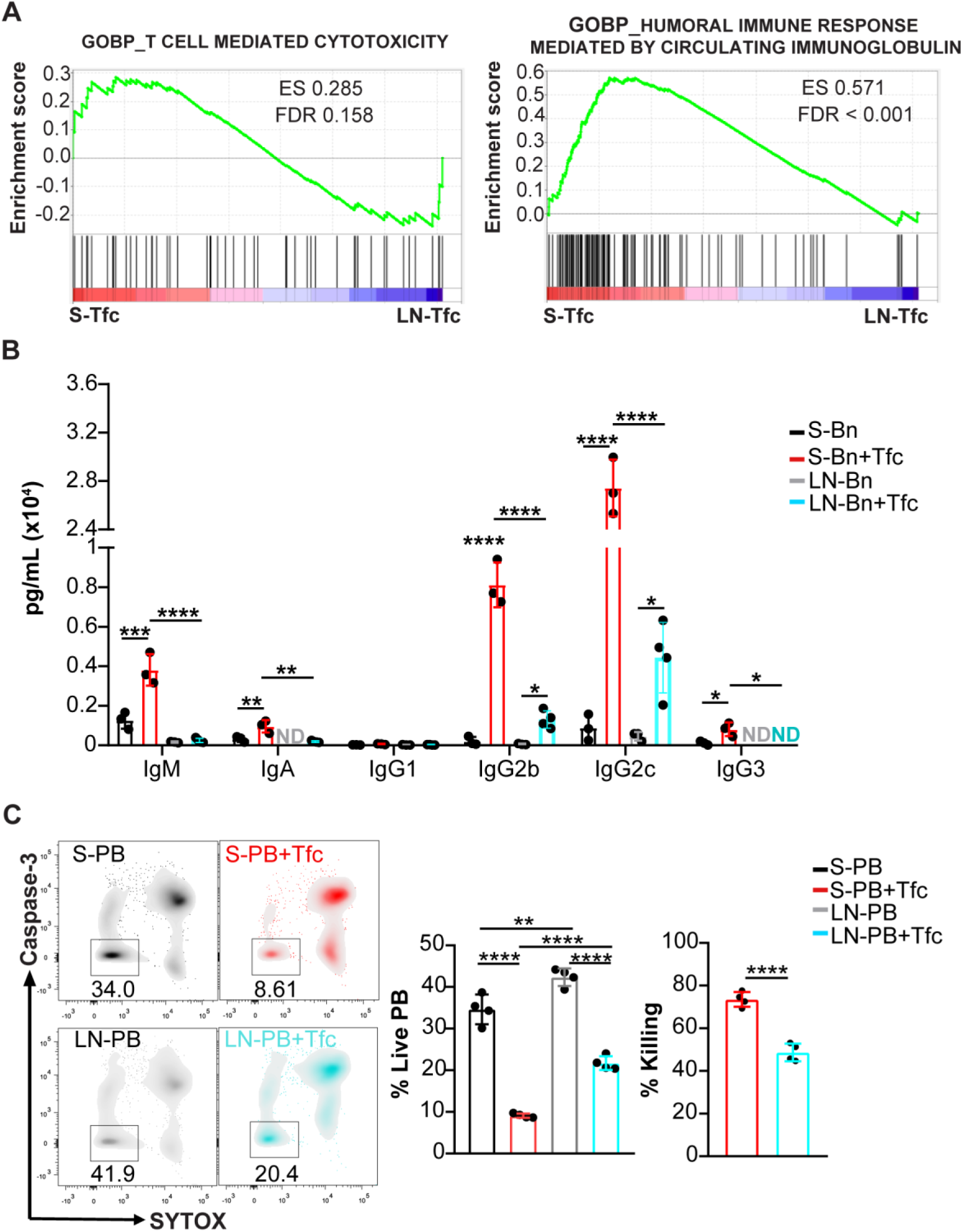
Analysis of helper and cytotoxic functions of S-Tfc and LN-Tfc cells. Gene Set Enrichment Analysis (GSEA) of bulk RNA-seq data from sorted splenic (S-Tfc, red) and inguinal LN-derived Tfc cells (LN-Tfc, cyan) obtained at 18 dpi, showing enrichment of *T cell-mediated cytotoxicity* and *humoral immune response mediated by circulating immunoglobulin* pathways (GO Biological Process, MSigDB mouse collection). ES, enrichment score; FDR, false discovery rate. Genes enriched in S-Tfc cells are located toward the left of the ranked list, whereas genes preferentially expressed in LN-Tfc cells are located toward the right. (B) Sorted naïve B cells (Bn) or (C) plasmablasts (PB) from spleen or inguinal LN were cultured for 20 h either alone (S-Bn/S-PB, black; LN-Bn/LN-PB, gray) or in the presence of sorted S-Tfc (red) or LN-Tfc (cyan) cells. All cell populations were purified from *T. cruzi*–infected mice at 18 dpi. (B) Statistical analysis of immunoglobulin concentrations (IgM, IgA, IgG1, IgG2b, IgG2c, and IgG3) measured in the culture supernatants using a multiplex bead-based assay (LEGENDplex). (C) Statistical analysis of the frequency of live plasmablasts (Caspase-3^-^SYTOX^-^CD8^-^B220^int^) and percentage of killing. ND, not detected. Each dot represents an individual mouse. Data are representative of three independent experiments (B, C). Statistical significance was determined using ordinary one-way ANOVA with selected comparisons and Bonferroni correction (*p < 0.05, **p < 0.01, ***p < 0.001, ****p < 0.0001).

Analysis of supernatants from naïve B cell co-cultures with Tfc cells derived from either SLO revealed that LN-Tfc cells significantly increased IgG2b and IgG2c production compared with B cells cultured alone. However, co-cultures containing S-Tfc cells resulted in significantly higher Ig levels overall and promoted a broader isotype distribution, including IgM, IgA, IgG2b, IgG2c, and IgG3. Notably, IgG3 was undetectable in supernatants from LN-Tfc co-cultures (Figure 6B).

We next examined the effect of Tfc cells on plasmablast survival. Co-culture of plasmablasts with Tfc cells from each SLO resulted in a significant reduction in the frequency of live plasmablasts compared with plasmablasts cultured alone (Figure 6C). Plasmablast viability was normalized to the corresponding tissue and expressed as percent killing, enabling direct comparison between SLO. Under these conditions, co-culture with S-Tfc cells resulted in a greater reduction in plasmablast viability compared with co-culture with LN-Tfc cells (Figure 6C).

## Discussion

In this study, we demonstrate that CD8^+^ Tfc cells emerging during the acute phase of *T. cruzi* infection share the canonical CXCR5^+^PD-1^+^CD8^+^CD3^+^ phenotype and exhibit comparable kinetics and parasite antigen specificity in the spleen and inguinal LN, yet diverge in their transcriptional, metabolic, phenotypic, and functional programs according to their SLO of residence. Together, these findings indicate that spatial context within SLO is associated with distinct patterns of Tfc cell differentiation and activity during infection.

Bulk RNA-seq analysis revealed that transcriptional segregation was primarily driven by tissue origin, as both Tfc and non-Tfc CD8^+^ T cells clustered according to the SLO analyzed. This indicates that CD8^+^ T cell transcriptional programs differ across SLO during *T. cruzi* infection. Similar tissue-associated specialization has been reported for other immune cell populations. Comparisons between plasma cells from the small intestine lamina propria and those residing in the bone marrow have demonstrated distinct transcriptional programs (30). Likewise, studies on innate lymphoid cells have shown that immune cell phenotypes and functions vary across lymphoid tissues in response to local environmental cues (31). Our findings extend these observations to Tfc cells during *T. cruzi* infection.

The transcriptional divergence observed between S-Tfc and LN-Tfc cells was accompanied by distinct metabolic features. S-Tfc cells displayed enrichment of glycolysis- and mTORC1-associated gene sets, increased glucose uptake, higher expression of GLUT1 and MCT1, and enhanced phosphorylation of p70 S6K, consistent with an effector-oriented metabolic configuration (25,32,33). They also exhibited increased mitochondrial mass and elevated mROS production, features commonly associated with activated CD8^+^ T cells (27,34). In contrast, LN-Tfc cells showed increased phosphorylation of AKT at Ser473, suggestive of enhanced mTORC2-associated signaling (25). The mTOR pathway integrates nutrient availability and signaling cues to coordinate T cell metabolic programs. In CD8^+^ T cells, mTORC1 activity has been linked to effector differentiation, whereas mTORC2 signaling has been associated with memory cell development and maintenance (35). These observations align with reports showing that immune cell function can differ across anatomical sites in association with distinct metabolic configurations, as described for gut- and bone marrow–resident IgA^+^ plasma cells exhibiting divergent bioenergetic profiles (36). In line with these metabolic features, Tfc subset composition also differed across SLO. S-Tfc cells were enriched in Eff/Em (CD44^+^CD62L^-^) populations, whereas LN-Tfc cells displayed a more heterogeneous distribution, with increased CM and memory precursor subsets. In agreement with this phenotype, LN-Tfc cells expressed higher levels of CD122 and CD127, receptors associated with IL-2– and IL-7–mediated survival and maintenance of memory CD8^+^ T cells (28,37). LN-Tfc cells also exhibited higher expression of Eomes, a transcription factor linked to memory-associated CD8^+^ T cell programs and long-term maintenance (38), together with T-bet expression (39), suggesting a mixed differentiation profile rather than terminal effector commitment. Notably, Tfc cells were preferentially detected in LN at later stages of infection (60 and 150 dpi), whereas they were nearly absent from the spleen, indicating differential persistence across SLO. In line with these observations, Tsui et al. (40) reported that LN harbor more heterogeneous virus-specific CD8^+^ T cell populations, whereas splenic cells are predominantly effector-like, supporting the concept that distinct SLO are associated with different CD8^+^ T cell states during infection.

Functionally, Tfc cells from both SLO retained the dual helper and cytotoxic capacities previously attributed to this subset (5,41,42). However, quantitative differences were evident. S-Tfc cells exhibited higher frequencies of cytotoxic mediators, including CD107a, Prf, and GrzB, and produced higher levels of IL-6 and IL-10 under polyclonal stimulation, whereas LN-Tfc cells produced higher levels of IL-2, consistent with their enrichment in memory-associated subsets. In co-culture assays, S-Tfc cells promoted broader immunoglobulin production, including IgM, IgA, IgG2b, IgG2c, and IgG3, and induced greater plasmablast killing compared with LN-Tfc cells. Although the precise mechanisms underlying these differences remain to be fully elucidated, enhanced CD40L and FasL expression on S-Tfc cells may contribute to increased immunoglobulin production and plasmablast cytotoxicity (5,43–45).

Together, these findings provide a comparative characterization of Tfc cells across spleen and LN during *T. cruzi* infection, revealing coordinated differences in transcriptional programs, metabolic configuration, differentiation state, functional output, and persistence according to anatomical location. These results underscore how distinct lymphoid tissue environments are associated with divergent CD8^+^ T cell specialization during infection and establish a foundation for future studies aimed at defining the metabolic and molecular mechanisms underlying Tfc cell heterogeneity.

## Author Contributions

Y.G. designed and performed the experiments, analysed the data, generated the figures, and wrote the original draft of the manuscript. A.G. and E.V.A.R. designed and supervised the study, contributed to methodological development, secured funding, and critically revised the manuscript. L.A. and J.C.G. contributed to sample collection and processing and participated in manuscript revision. C.C.S. provided guidance in metabolic assay design and analysis and revised the manuscript. C.L.M. contributed to project administration, data analysis, and manuscript revision. RNA sequencing was performed by AZENTA. All authors contributed to the article, approved the submitted version, and agree to be accountable for all aspects of the work.

## Acknowledgements

We thank the staff of the facilities at CIBICI: M. P. Abadie, M. P. Crespo, S. Boccardo, V. Blanco, D. Lutti, C. Noriega, F. A. Frontera, S. R. Oms, R. E. Villarreal, G. Furlán, N. M. Maldonado, M. S. Miró for their excellent technical assistance. We acknowledge the NIH Tetramer Core Facility for provision of Brilliant Violet 421-labeled TSKB20 tetramers.

## Ethics statement

All procedures were approved and conducted in accordance with the guidelines of the Institutional Animal Care and Use Committee of the Facultad de Ciencias Químicas, Universidad Nacional de Córdoba (protocol numbers RD-723 2022 and RD-349–2022).

## Data Availability Statement

RNA-seq data are publicly available in the NIH Sequence Read Archive (PRJNA1234210, https://www.ncbi.nlm.nih.gov/sra/PRJNA1234210). The data discussed in this publication have been deposited in NCBI’s Gene Expression Omnibus (Edgar *et al*., 2002) and are accessible through GEO Series accession number GSE319904 (https://www.ncbi.nlm.nih.gov/geo/query/acc.cgi?acc=GSE319904). Other supporting data are available from the corresponding author upon reasonable request.

## Supporting information

**Figure S1.** Transcriptional analysis of Tfc and non-Tfc cells from spleen and LN from *T. cruzi* infected mice. CXCR5^+^PD-1^+^ (Tfc) and CXCR5^-^PD-1^-^ (non-Tfc) CD8^+^ T cells were isolated by cell sorting from spleen and inguinal LN of *T. cruzi*–infected mice at 18 dpi and subjected to bulk RNA sequencing. Principal component analysis of the whole transcriptome shows the distribution of Tfc and non-Tfc samples from both SLO. Each dot represents one biological replicate (*N* = 3), consisting of pooled cells from three mice.

**Table S1.** Abs and reagents used for flow cytometry. Detailed list of Abs and reagents used in flow cytometry experiments, including their target specificity, fluorochrome conjugate, supplier, catalog number, clone, and working dilution.

