## Supplementary Figure 1 and Table 1 for "Tissue-distinct Features of Follicular Cytotoxic CD8^+^ T Cells in *Trypanosoma cruzi* infection": Table S1.pdf

| Surface markers | | | | | |
|---|---|---|---|---|---|
| **Target** | **Fluorochrome/Biotin** | **Company** | **Catalog #** | **Clone** | **Dilution** |
| B220 | FITC | Biolegend | 103202 | RA3-6B2 | 1/200 |
| B220 | PerCp-eFluor.710 | eBioscience | 46-0452-82 | RA3-6B2 | 1/200 |
| B220 | PE-Cy7 | eBioscience | 25-0452-82 | RA3-6B2 | 1/300 |
| B220 | BV421 | Biolegend | 103240 | RA3-6B2 | 1/400 |
| B220 | APC-Cy7 | Biolegend | 103224 | RA3-6B2 | 1/200 |
| B220 | BV711 | Biolegend | 103255 | RA3-6B2 | 1/300 |
| CD122 | PE-Cy7 | eBioscience | 25-1222-80 | TM-b1 | 1/100 |
| CD127 | BV711 | Biolegend | 135035 | A7R34 | 1/100 |
| CD138 | APC | BD Biosciences | 561705 | 281-2 | 1/200 |
| CD138 | BV421 | BD Biosciences | 562610 | 281-2 | 1/200 |
| CXCR5 | Biotin | BD Pharmingen | 551960 | 2G8 | 1/75 |
| ICOS | PerCp-eFluor.710 | eBioscience | 46-9942-82 | 7E.17G9 | 1/100 |
| CD3e | PerCp/Cy5.5 | eBioscience | 45-0031-82 | 145-2c11 | 1/100 |
| CD4 | FITC | eBioscience | 11-0041-85 | GK1.5 | 1/200 |
| CD4 | PE-Cy7 | eBioscience | 25-0041-82 | GK1.5 | 1/200 |
| CD4 | APC | Biolegend | 100530 | GK1.5 | 1/200 |
| CD4 | APC-eFluor 780 | eBioscience | 47-0041-82 | GK1.5 | 1/200 |
| CD4 | PECF594 | BD Biosciences | 562285 | RM4-5 | 1/300 |
| CD40L | PE | BD Pharmingen | 553658 | MR1 | 1/100 |
| CD44 | PE-Cy7 | Biolegend | 103030 | IM7 | 1/200 |
| CD44 | PE-Cy5 | eBioscience | 15-0441-81 | IM7 | 1/200 |
| CD62L | APC | eBioscience | 17-0621-83 | MEL-14 | 1/200 |
| CD62L | SB 600 | eBioscience | 63-0621-82 | MEL-14 | 1/200 |
| CD8a | FITC | eBioscience | 11-0081-86 | 53-6.7 | 1/200 |
| CD8a | AlexaFluor 700 | eBioscience | 56-0081-82 | 53-6.7 | 1/200 |
| CD8a | PE-Cy7 | eBioscience | 25-0081-82 | 53-6.7 | 1/300 |
| FasL | PE | BD Biosciences | 555293 | MFL3 | 1/100 |
| GLUT1 | Biotin | Novus Biologicals | NB110-39113B | Polyclonal | 1/100 |
| H-2Kb/TSKB20 | BV421 | NIH Tetramer Core Facility | - | - | 1/400 |
| IgD | FITC | eBioscience | 11-5993-82 | 11-26c | 1/200 |
| KLRG1 | APC | Biolegend | 138412 | 2F1/KLRG1 | 1/200 |
| MCT1 (SLC16A1) | APC | eBioscience | AMT-011-APC50UL | Polyclonal | 1/100 |
| PD-1 | PE-Cy7 | eBioscience | 25-9985-82 | J43 | 1/100 |
| PD-1 | APC | eBioscience | 17-9985-82 | J43 | 1/100 |
| PD-1 | Brillant Violet 421 | Biolegend | 135221 | 29F.1A12 | 1/100 |

| Secondary reagents | | | | | |
|---|---|---|---|---|---|
| **Reagent** | **Fluorochrome** | **Company** | **Catalog #** | **Clone** | **Dilution** |
| Goat anti-Rabbit IgG | Alexa Fluor 546 | eBioscience | A-11071 | Polyclonal | 1/500 |
| Streptavidin | PE | eBiosciences | 12-4317-87 | - | 1/300 |
| Streptavidin | PE-Cy7 | eBioscience | 25-4317-82 | - | 1/300 |
| Streptavidin | APC | BD | 554067 | - | 1/300 |

| Intracellular / transcription factors / cytokines | | | | | |
|---|---|---|---|---|---|
| **Target** | **Fluorochrome** | **Company** | **Catalog #** | **Clone** | **Dilution** |
| Bcl-6 | AlexaFluor 647 | BD Bioscience | 563582 | K112-91 | 1/75 |
| CD107a | PE | Biolegend | 121612 | 1D4B | 1/200 |
| CD107a | APC/Cy7 | eBioscience | 121616 | 1D4B | 1/150 |
| Eomes | PE-eFluor610 | eBioscience | 61-4875-82 | Dan11mag | 1/400 |
| Granzyme A | PerCp-eFluor.710 | eBioscience | 46-5831-80 | GzA-3G8.5 | 1/200 |
| Granzyme B | FITC | Biolegend | 515403 | GB11 | 1/100 |
| IFNg | BV711 | BD Bioscience | 564336 | XMG1.2 | 1/200 |
| IL-21 | APC | eBioscience | 17-7211-82 | FFA21 | 1/50 |
| IL-10 | PE | Biolegend | 505008 | Jes5-16E3 | 1/100 |
| IL-6 | eFluor450 | eBioscience | 48-7061-82 | MP5-20F3 | 1/75 |
| IL-2 | BV421 | Biolegend | 503837 | JES6-5H4 | 1/75 |
| IRF4 | PerCp-eFluor.710 | eBioscience | 46-9858-82 | 3E4 | 1/100 |
| Perforin | APC | Biolegend | 154304 | S16009A | 1/100 |
| Phospho-Akt (Ser473) | PE | Cell Signaling | 4060 s | D9E | 1/100 |
| Phospho-mTOR (Ser2448) | PerCp-eFluor.710 | eBioscience | 46-9718-42 | MRRBY | 1/100 |
| Phospho-p70 S6 Kinase (Thr389) | - | Cell Signaling | 9234T | 108D2 | 1/75 |
| T-bet | PE-Cy7 | eBioscience | 25-5825-82 | 4B10 | 1/100 |
| TCF-1 | PE | BD Bioscience | 564217 | S33-966 | 1/200 |
| TNF | eFluor450 | eBioscience | 48-7321-82 | MP6-XT22 | 1/200 |

| Metabolic / functional probes | | | |
|---|---|---|---|
| **Reagent** | **Company** | **Catalog #** | **Concentration** |
| Mito Tracker Green FM | ThermoFisher Scientific | M-7514 | 100 nM |
| MitoTracker Orange CMTMRos | ThermoFisher Scientific | M-7510 | 100 nM |
| MitoStatus™ Red | BD | 564697 | 100 nM |
| MitoSOX Red | ThermoFisher | M36008 | 5 uM |
| 2-NBDG | ThermoFisher | N13195 | 10 nM |

| Viability Dyes | | | |
|---|---|---|---|
| **Reagent** | **Company** | **Catalog #** | **Dilution** |
| Live/Dead Fixable Aqua 405 | eBioscience | L34966 | 1/400 |
| Live/Dead NIR Fixable Viability Kit 633 | eBioscience | L10119 | 1/800 |
| CellEvent™ Caspase-3/7 Green | eBioscience | C10427 | |

### Table S1
