## Supplementary figures and images for "Tissue-distinct Features of Follicular Cytotoxic CD8^+^ T Cells in *Trypanosoma cruzi* infection"

### Figure S1.tif

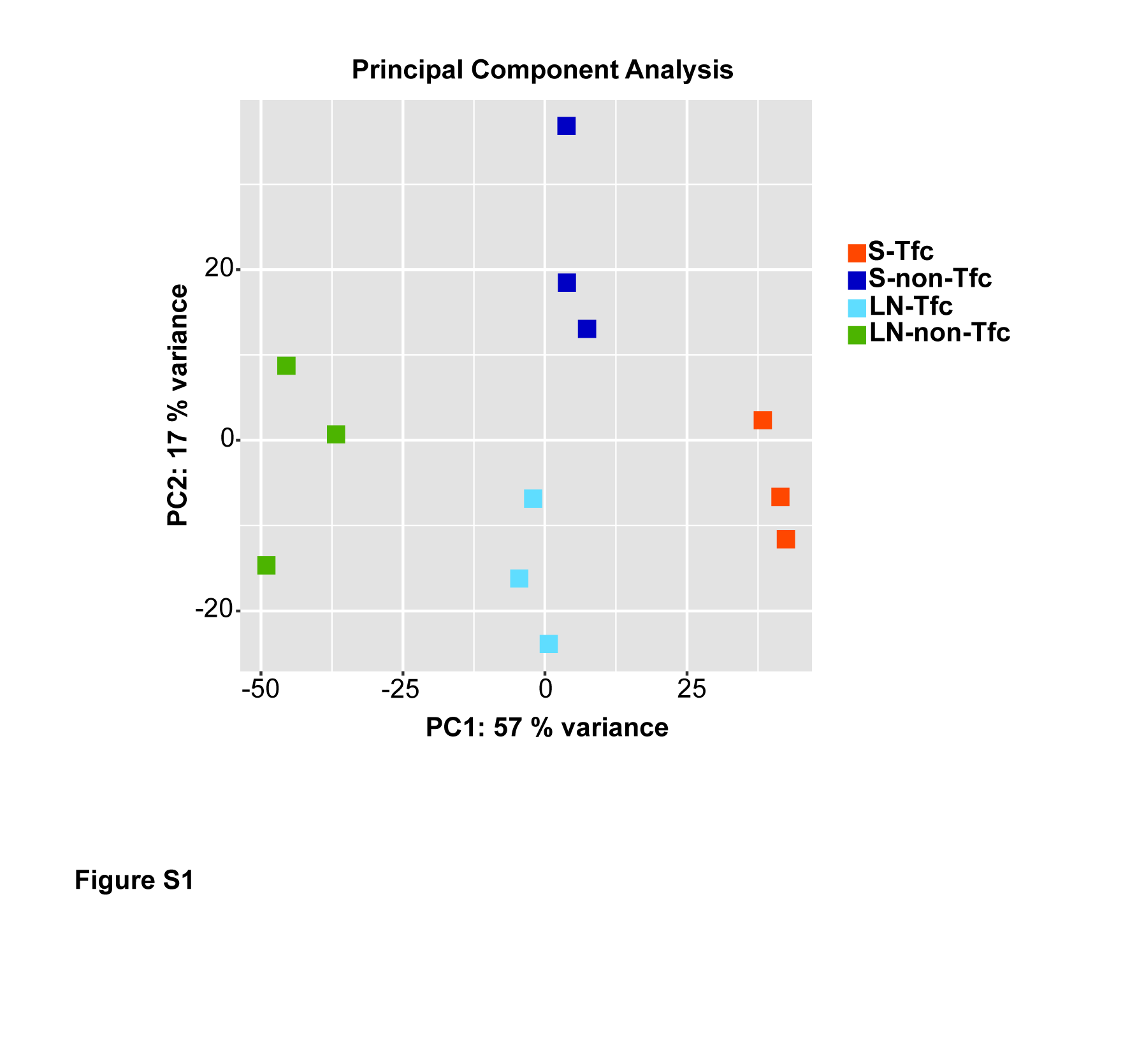
